# Tangled genetic relationships within the *Fusobacterium* genus

**DOI:** 10.1101/2024.03.27.587012

**Authors:** Cristian Molteni, Diego Forni, Rachele Cagliani, Manuela Sironi

**Affiliations:** Scientific Institute IRCCS E. MEDEA, Bioinformatics, 23842 Bosisio Parini, Italy

## Abstract

Fusobacteria have been associated to different diseases, including colorectal cancer (CRC), but knowledge of which taxonomic groups contribute to specific conditions is incomplete. We analyzed the genetic diversity and relationships within the *Fusobacterium* genus. We report recent and ancestral recombination in core genes, indicating that fusobacteria have mosaic genomes and emphasizing that taxonomic demarcation should not rely on single genes/gene regions. Across databases, we found ample evidence of species miss-classification and of undescribed species, which are both expected to complicate disease association. By focusing on a lineage that includes *F. periodonticum/pseudoperiodonticum* and *F. nucleatum*, we show that genomes belong to four modern populations, but most known species/subspecies emerged from individual ancestral populations. Of these, the *F. periodonticum/pseudoperiodonticum* population experienced the lowest drift and displays the highest genetic diversity, in line with the less specialized distribution of these bacteria in oral sites. A highly drifted ancestral population instead contributed genetic ancestry to a new species, which includes genomes classified within the *F. nucleatum animalis* diversity in a recent CRC study. Thus, evidence herein calls for a re-analysis of *F. nucleatum animalis* features associated to CRC. More generally, our data inform future molecular profiling approaches to investigate the epidemiology of *Fusobacterium*-associated diseases.

## Introduction

Fusobacteria are Gram-negative, non-spore-forming obligate anaerobes with a wide distribution. The phylum Fusobacteriota includes both species commonly found in animal microbiota and others that are free-living in the marine environment^1^. Within the phylum, members of the genus *Fusobacterium* are found in the mouth and other mucosal sites of humans and other animals^1^. In the human oral cavity, *Fusobacterium* species participate to the formation of polymicrobial biofilms and can cause periodontal disease. Remarkably, these bacteria have the ability to spread to extraoral sites where they contribute to the development of different conditions, including Lemierre syndrome, appendicitis, brain abscesses, osteomyelitis, pericarditis, adverse pregnancy outcomes, inflammatory bowel disease, and cancer^1^. *Fusobacterium* species have gained enormous interest in relation to their oncogenic role in colorectal cancer (CRC) and other tumor types. In particular, most studies have focused on *Fusobacterium nucleatum*, which was shown to be enriched in the gut microbiota of CRC patients and to promote carcinogenesis through multiple mechanisms^1–6^. *F. nucleatum (*and *F. necrophorum)* was also found in the lymph node and liver metastases of *Fusobacterium*-associated primary tumors^3^. Growing evidence however suggests that *Fusobacterium* species other than *F. nucleatum* associate with CRC^3,7–10^. Moreover, *F. nucleatum* bacteria are presently classified into four subspecies (*nucleatum*, *animalis*, *vincentii,* and *polymorphum).* These subspecies are phylogenetically divergent to the point that they were suggested to represent distinct species^11,12^. Also, a recent report suggested that *F. nucleatum* subspecies *animalis* is divided into two clades, only one of which is associated with CRC^6^. Finally, several works identified species miss-classifications in public records, whereas some *Fusobacterium* genomes cannot be assigned to any existing species^9,13^.

Recently, it was suggested that phylogenetic analyses based on the *rpoB* gene, rather than on 16s rRNA, are better suited to differentiate *Fusobacterium* species and to classify genomes into lineages^9^. However, analyses in several human commensal microbiota and environmental bacteria have suggested that homologous recombination may affect the majority of loci in the genome^14–20^. Thus, for many species, each locus has recombined extensively, and consequently the phylogeny changes many times along the genome alignment, making it impossible to reconstruct robust clonal relationships. This might also be the case of fusobacteria, as a study limited to *F. necrophorum* revealed extensive recombination^21^. Nonetheless, the extent of recombination in the extended *Fusobacterium* genus has not been investigated, making it difficult to assess how well single gene-based phylogenies can represent the relationships among genomes. Also, a comprehensive analysis of the genetic diversity and of evolutionary relationships in this genus is presently missing.

## Results

### Recombination in *rpoB* and relevance for lineage definition

The *rpoB* gene was recently suggested to represent a good marker for the classification and the phylogenetic reconstruction of relationships among members of the genus *Fusobacterium*^9^. We thus retrieved from public databases sequence information for 361 *Fusobacterium* genomes and we extracted *rpoB* sequences, which were identified for 345 strains. The neighbor-net split network of *rpoB* showed a complex reticulation pattern, suggesting extensive recombination (Figure 1). In line with previous reports, *F. naviforme* sequences, as well as some other unassigned species, were highly divergent^7,9,22^. Given the observed reticulation, we used the fastGEAR software to identify and analyze recombination events. This software first classifies the sequences into lineages; subsequently, it calculates the number of ancestral and recent recombination events and tests for their significance. fastGEAR divided *rpoB* sequences into 12 lineages and identified 198 ancestral recombination events and 98 recent ones (Figure 1, Supplementary Figure S1, Supplementary Table 1). This clearly indicates the presence of extensive recombination and implies that different gene regions have distinct evolutionary histories. We thus used SimPlot analysis to generate a sequence similarity network, which joins nodes (sequences or groups of sequences) with edges when similarity is above a given threshold. With a threshold for global and local similarities at 95%, the twelve lineages remained separated (Figure 2A). However, several regions of local similarity above 95% were detected, in line with the effects of recombination.

**Figure 1.**
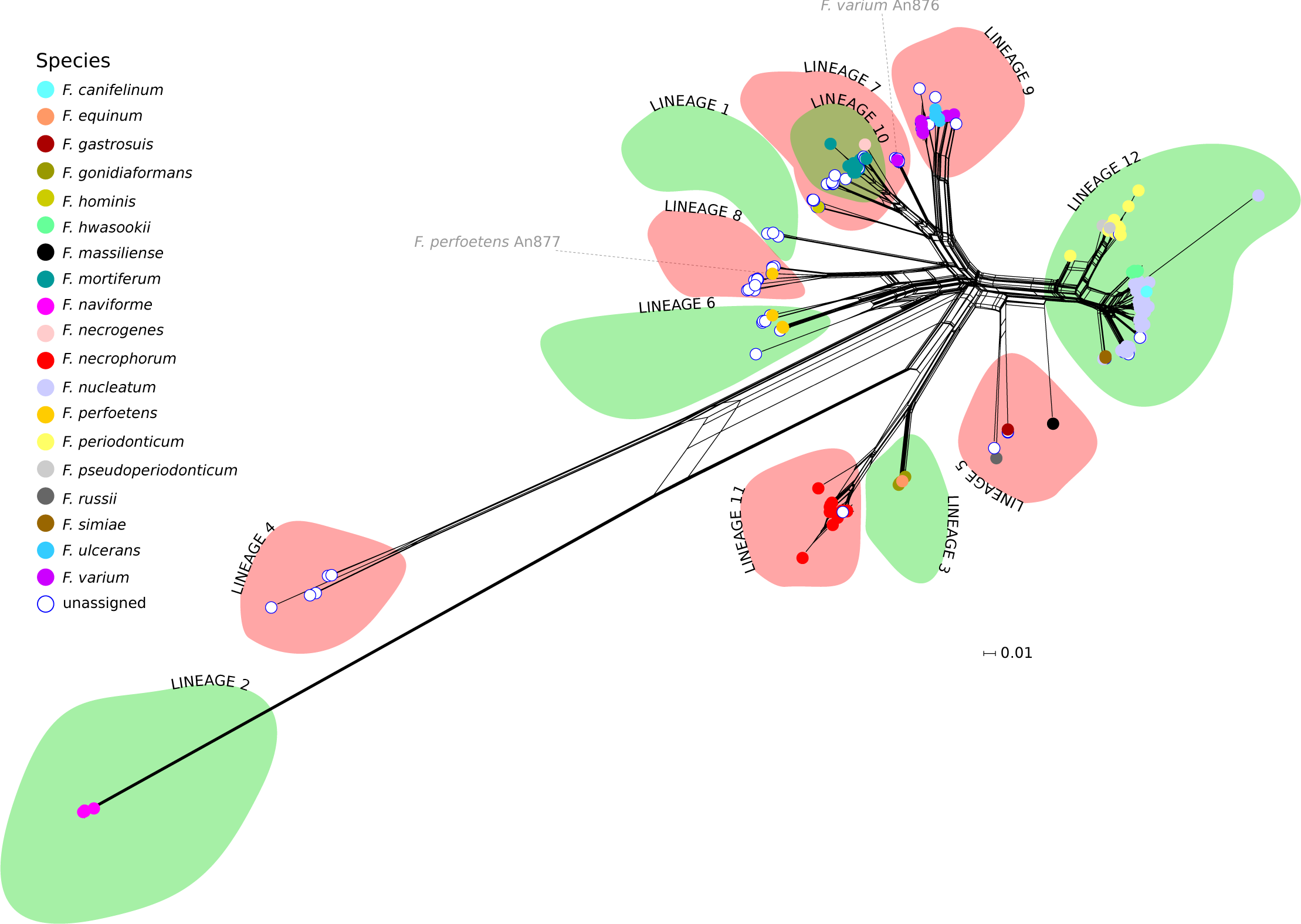
Recombination and divergence among *Fusobacterium* species. **(A)** Neighbor-net split network of 345 *rpoB* genes. Each sequence is shown as a dot, color-coded by species. The green and red areas represent the lineages defined by fastGEAR analysis (see Methods). The *F. varium* An876 and *F. perfoetens* An877 sequences are highlighted in gray (see also Figure 2B and Figure 4).

**Figure 2.**
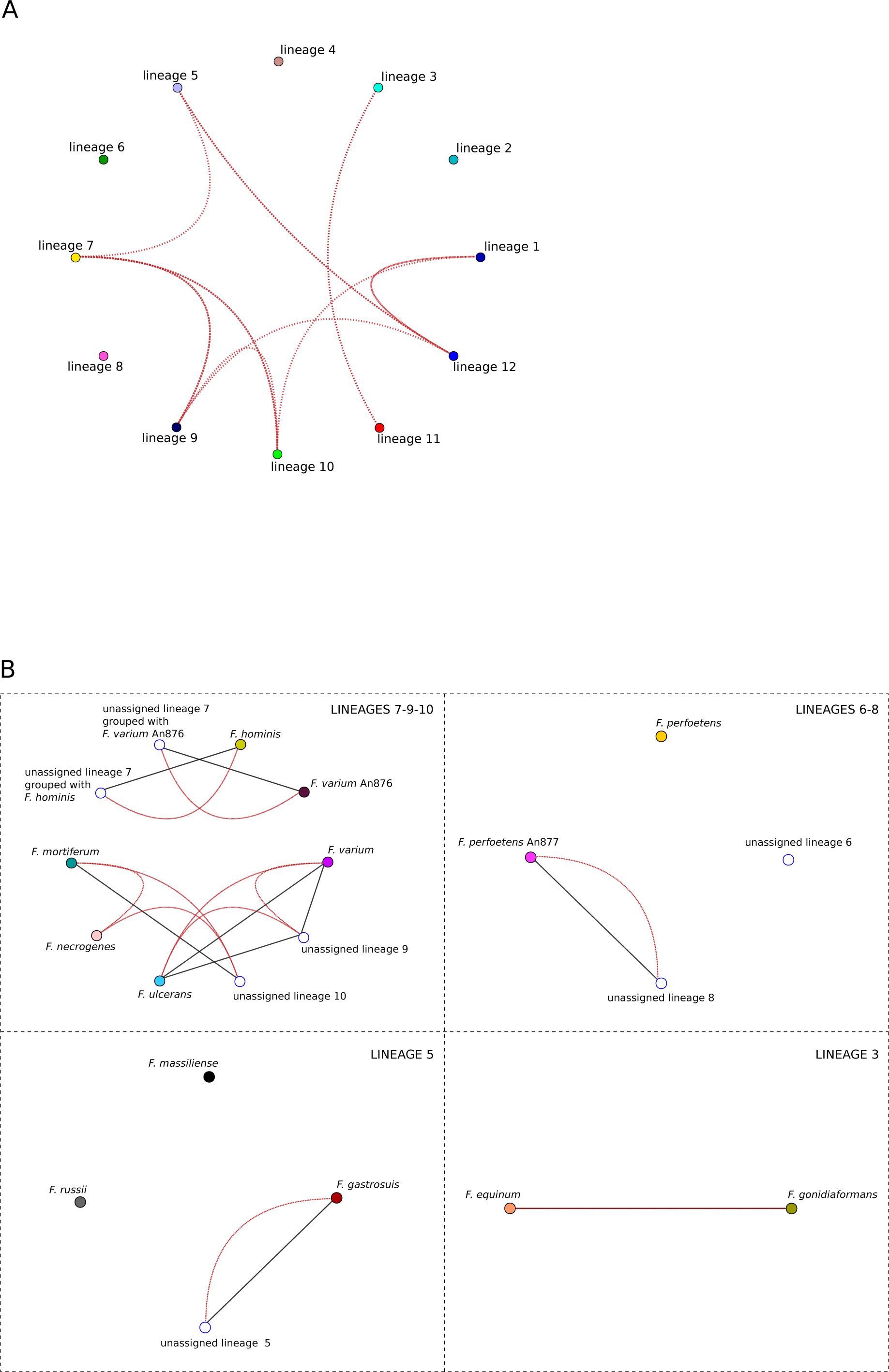
Sequence similarities among lineages. **(A)** Sequence similarity network based on *rpoB* genes. Sequences are grouped by lineages, as defined by fastGEAR. Global sequence similarity is represented by black edges and it is not observed among rpoB sequences. Red edges represent local similarity. Thresholds for global and local similarities were set to 95%. **(B)** Sequence similarity networks based on *rpoB* sequences are shown for selected lineages (see main text). As in panel A, lineages are color-coded as in Figure 1. Thresholds for global and local similarities were set to 95%.

On one hand, the lineage subdivision generated by fastGEAR showed good agreement with the clusters in the neighbor-net split network, with the exclusion of lineage 7, which was split into two clusters (Figure 1). On the other, the identified lineages only partially reflected the demarcation of known *Fusobacterium* species and two lineages (1 and 4) were only populated with unclassified fusobacteria. In several instances, more than one species was classified in the same lineage, whereas in the case of *F. varium* and *F. perfoetens*, genomes were split into two different lineages (Figure 1). Specifically, one *F. varium* sequence (strain An876) was assigned to lineage 7 together with some unassigned species and with *F. hominis*, while all the others were in lineage 9, which also includes *F. ulcerans*. Likewise, one *F. perfoetens rpoB* sequence (strain An877) and several unassigned species were in lineage 8, whereas the remaining ones were assigned to lineage 6. Finally, multiple species were detected in lineages 3 (*F. equinum* and *F. gonidiaformans*) and 5 (*F. russii, F. massiliense, F. gastrosuis*) (Figure 1).

We thus used SimPlot to analyze the similarity between the *rpoB* sequences in these lineages. Briefly, results (Figure 2B) indicated that i) the F. varium sequence in lineage 7 shows less than 95% local and global similarity to other *F. varium* sequences in lineage 9 and the same holds true for the *F. perfoetens* sequence in lineage 8 compared to other *F. perfoetens* sequences; thus, these two sequences are likely to be misclassified; ii) *F. equinum* and *F. gonidiaformans* have high sequence similarity (>95%) (see below); iii) *F. russii, F. massiliense,* and *F. gastrosuis* display below threshold similarity at the global and local level. Overall, these data confirm previous indications that the *Fusobacterium* genus includes substantial unclassified diversity and that some sequences are miss-classified.

Finally, we compared the classification determined by fastGEAR with the nine lineages defined by Bi and coworkers^9^. This exercise was complicated by the fact that the analyzed sequences differ. Nonetheless, a relatively good correspondence was found. The main differences related to *F. necrophorum* and *F. gonidiaformans*, which fastGEAR classified in two distinct lineages, whereas they both contributed to the same lineage in Bi et al. On the contrary, fastGEAR classified *F. massiliense* and *F. russii* together whereas they accounted for lineages 2 and 3 in the previous classification proposed by Bi and coworkers^9^.

### Recombination in core genes and its effects on classification

We next aimed to investigate and compare the levels of recent and ancestral recombination among different core genes. We thus used the Genome Taxonomy Database Toolkit to extract the sequences of 120 core genes present in more than 300 fusobacterial genomes. Among these we retained only the ones longer than 1000 bp (n=45) and used fastGEAR to detect recombination. For these 45 genes, the inferred number of lineages varied from 8 to 19. In most cases, the levels of ancestral recombination were much higher than the recent. The number of ancestral events ranged from 37 to 446, whereas recent events ranged from 12 to 400 (Supplementary Table 2). Consistent with a relatively constant rate of recombination at individual genes, the number of ancient and recent events was highly correlated (Pearson’s correlation coefficient= 0.58, p value= 2.4×10^-05^). Because the amount of recombination events is clearly also a function of gene size, we normalized the number of events by alignment length, so as to have a measure of recombination intensity. The results indicated that *rpoB* is in the low range of recombination intensities, whereas the top recombining genes are involved in different functions such as DNA replication and repair (e.g., *polA* and *mfd*) and peptidoglycan biosynthesis (*murD*) (Figure 3).

**Figure 3.**
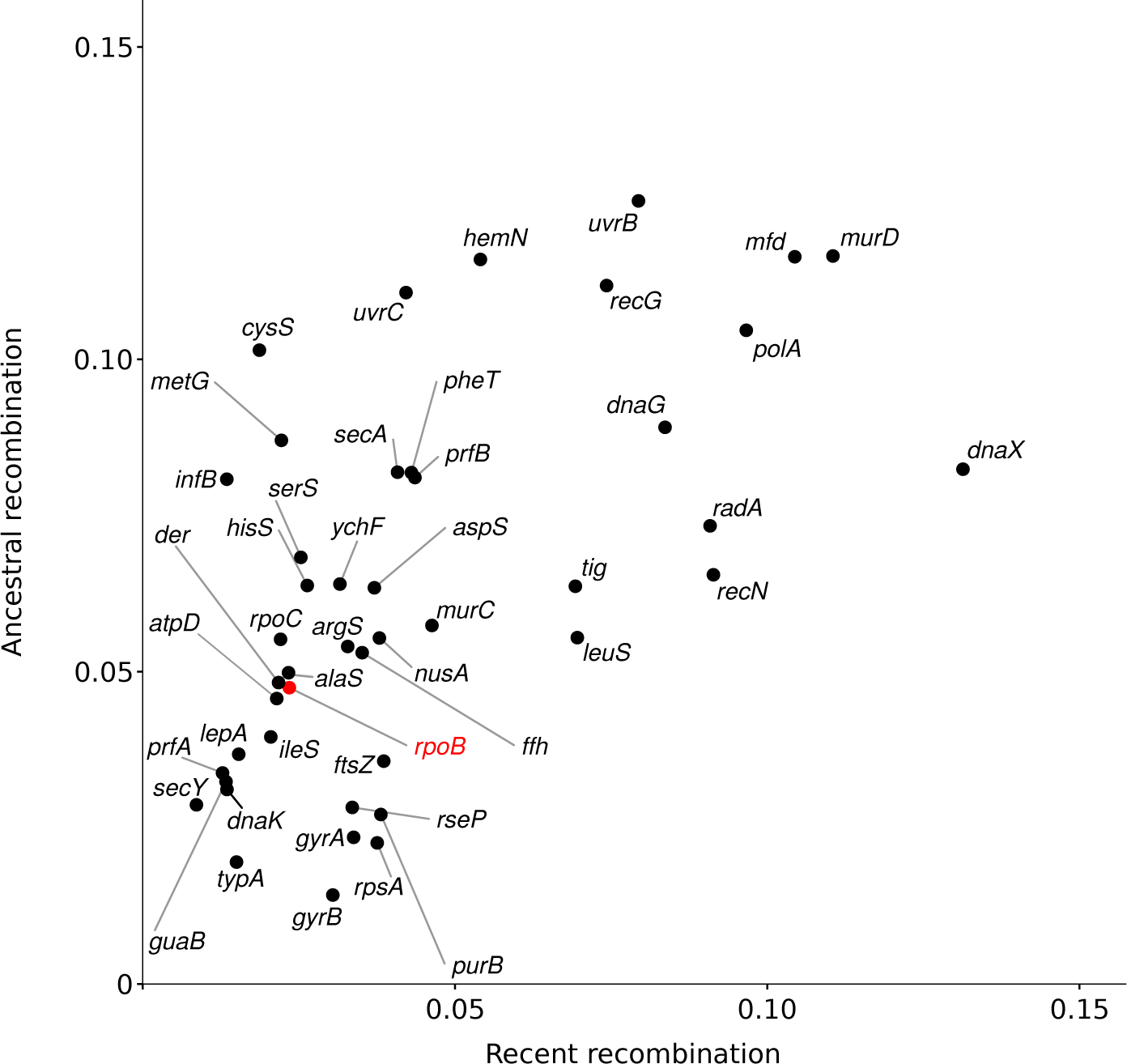
Recombination intensities in core genes. Correlation between ancestral and recent recombination events for 45 core genes. Recombination events were calculated with fastGEAR and normalized for alignment length. Each dot is labeled with the corresponding gene name and *rpoB* is highlight in red.

Overall, these results suggest that, in fusobacteria, individual genes have different evolutionary histories and recombination intensities. As a consequence, a classification based on individual genes is expected to be highly sensitive to the choice of the genomic region.

To gain further insight into the effect of gene choice, we again resolved to SimPlot analysis. In particular, we generated a concatenated alignment with the 120 core genes and we analyzed global similarity among species and lineages (Figure 4). Using a threshold of 95%, most edges joined nodes belonging to lineage 12 (*F. nucleatum*, *F. hwasookii*, *F. canifelinum* and *F. simiae*, as well as *F. periodonticum* and *F. pseudoperiodonticum*) and unassigned sequences therein. In line with a very recent report, *F. equinum* and *F. gonidiaformans* showed high similarity in the extended set of core genes, as well^22^ (Figure 1). We next compared global with local similarity patterns defined by three genes: *rpoB*, *typA* (with low recombination intensity) and *murD* (with high recombination intensity). In all cases local similarities were more extended than global similarity, indicating that classifications based on single genes tend to cluster together sequences that are divergent at the level of the extended set of core genes (Figure 4). Also, whereas the pattern of local similarity was relatively similar for the low recombining *rpoB* and *typA*, it was not for the highly recombining *murD* gene. Indeed, fewer cases of local high similarity were detected with *murD* and in some instances *murD* sequences were more divergent than 95% even between species that showed high global similarity (e.g., *F. simiae* and *F. nucleatum* or *F. hwasookii* and *F. canifelinum*) (Figure 4). These data underscore the effect of recombination on similarity scores and caution against using individual genes or gene regions for classification purposes.

**Figure 4.**
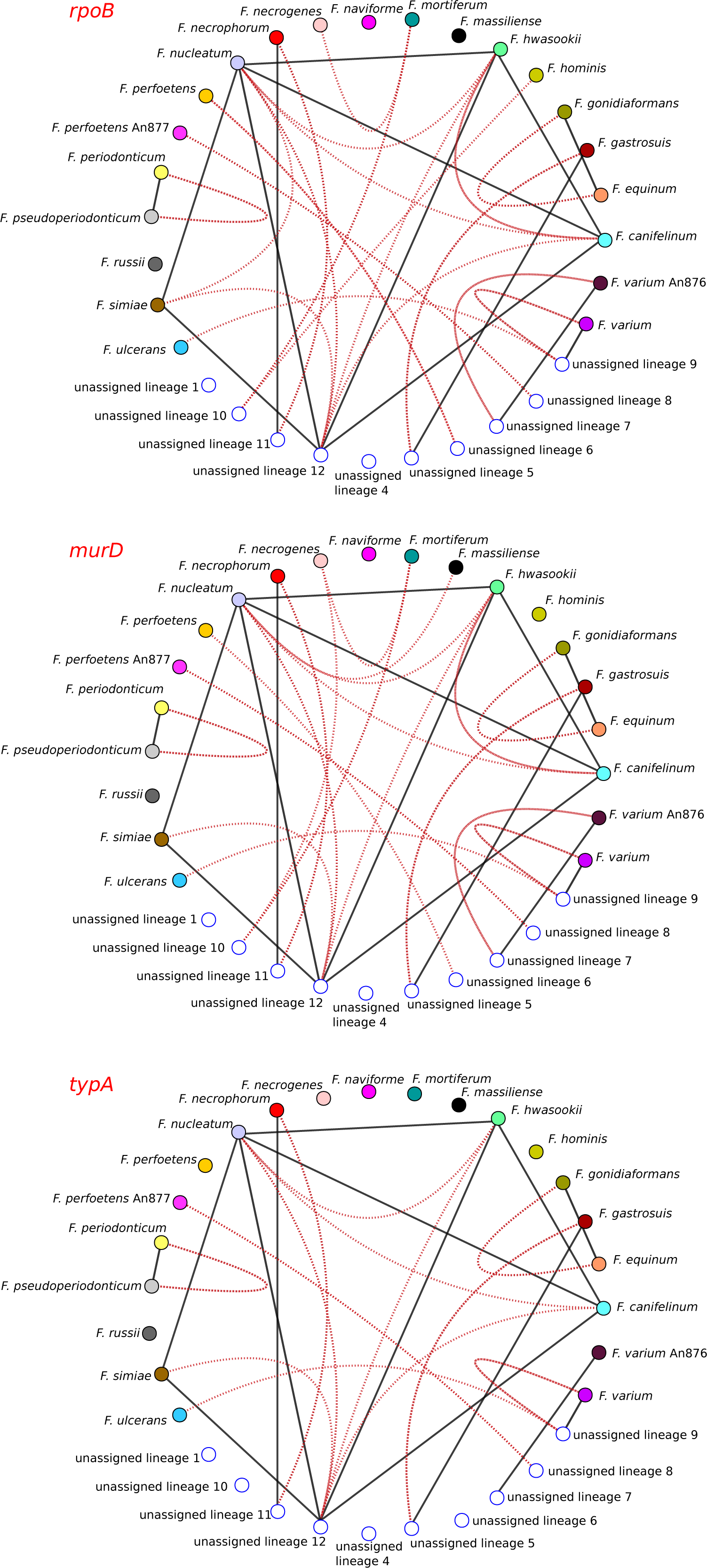
Sequence similarities among *Fusobacterium* species and lineages. Sequence similarity networks based on a concatenated core gene alignment. For all networks, global and local similarity thresholds were set to 95%. Black edges represent global sequence similarity (i.e. calculated for the whole concatenated gene alignment). Red edges display local similarity within three different selected genes in the alignment: *rpoB, murD,* and *typA.* Each species or lineage is shown as a colored node (colors as in Figure 1).

### Genetic relationships in the *F. nucleatum/F. periodonticum* lineage

We next aimed to investigate the genetic relatedness among fusobacteria in lineage 12 (lineage 1 in Bi and co-workers^9^) (Figure 1). This lineage comprises species often associated with CRC, including the highly studied *F. nucleatum* (and its subspecies), and the different species have high sequence similarity in the analysis of core genes (Figure 4). We thus extracted parsimony-informative (PI) sites from the core gene alignment. PI information was used as the input for principal component analysis (PCA). In agreement with the neighbor-net split tree, the first PC explained 32% of the variance and separated the two main sub-lineages – i.e., *F. periodonticum/pseudoperiodonticum* from the other *Fusobacterium* species (Figure 5A). Along this component, *F. periodonticum* separated into two sub-clusters, suggesting the presence of unrecognized diversity within this species. The second PC explained 19% of variance and separated the non-*F. periodonticum/pseudoperiodonticum* core genomes in three clusters: i) one containing *F. nucleatum polymorphum, F. hwasookii, F. canifelinum* and *F. nucleatum nucleatum,* plus several unclassified strains; ii) another comprising *F. nucleatum vincentii, F. simiae* and four unclassified genomes; iii) the third only featuring *F. nucleatum animalis* and unclassified species (Figure 5A). This clearly indicates that the four *F. nucleatum* subspecies are less closely related to each other than they are to other *Fusobacterium* species. Therefore, they should be considered as separate species.

**Figure 5.**
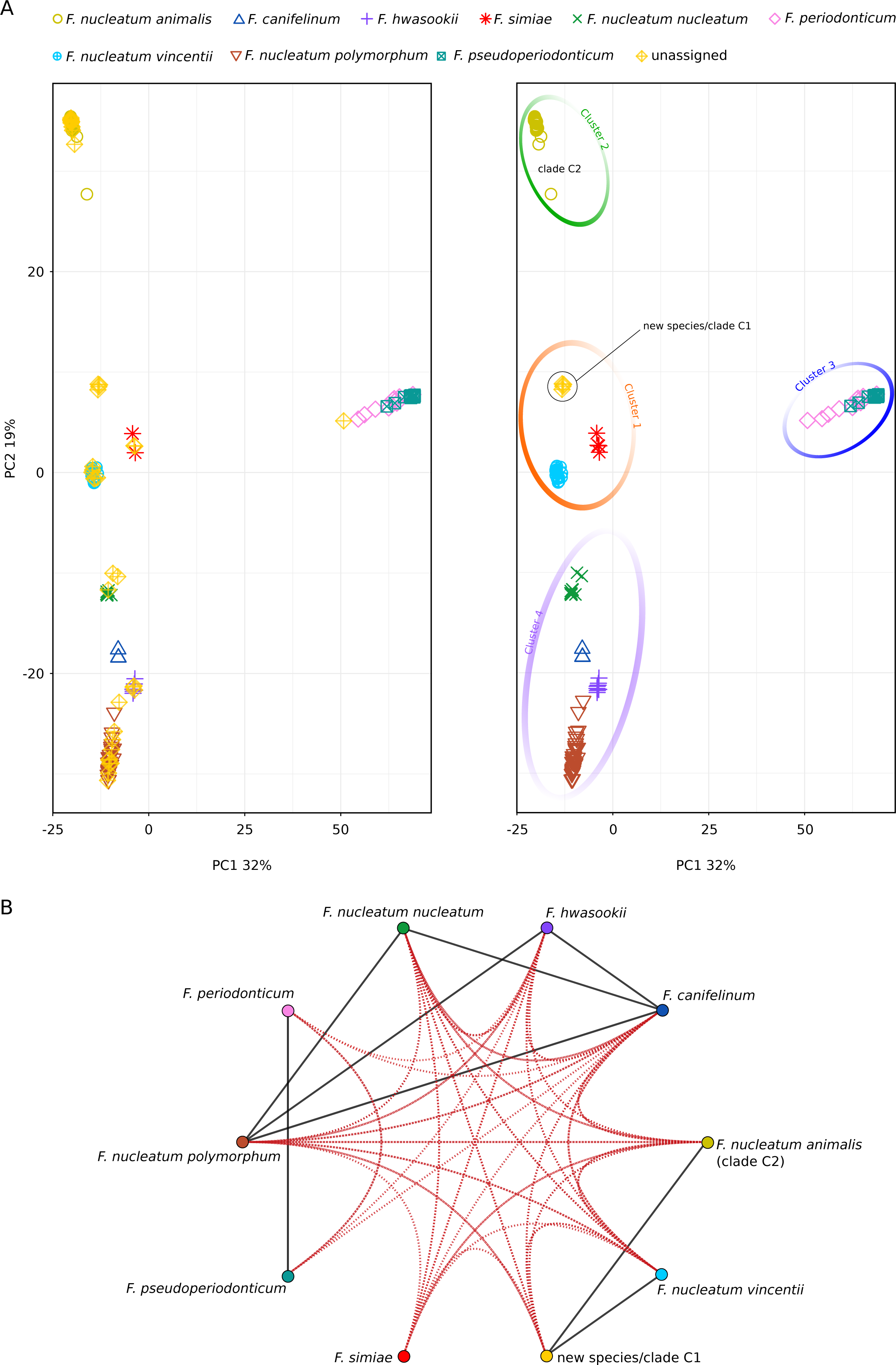
Principal Component Analysis (PCA) and similarity network for *Fusobacterium* genomes in the *F. nucleatum/F. periodonticum* lineage. **(A)**. Each *Fusobacterium* genome is colored and displayed with a different symbol, as described in the legend. On the left side, the plot shows several unassigned sequences, which were reassigned in the right panel. In the right panel, the four major clusters are highlighted by colored circles. The genomes contributing to the new species/clade C1 and clade C2 are indicated. **(B)** Sequence similarity network. Global and local sequence similarity are shown in black and red edges (thresholds were set to 95%). Nodes are colored as in panel A.

The results of the PCA were used to provisionally assign unclassified genomes to known species (Figure 5A). The only exception was accounted for by four unclassified genomes closely related to each other, suggesting they represent an undescribed species. Indeed, one of these is *Fusobacterium* FNU, previously suggested to represent a new species^13^. Another genome in this hypothetical new species belongs to strain 13-08-02 (BHYR00000000). Very recently, Zepeda-Rivera and coworkers reported that *F. nucleatum animalis* genomes can be divided into two clades referred to as C1 and C2, with the latter associated with the CRC niche^6^. Strain 13-08-02 was included in C1, whereas their clade C2 comprised a number of genomes classified as *F. nucleatum animalis* in NCBI and in the PCA analysis (Figure 5A). Overall, the PCA does not support the idea that genomes in clade C1 belong to *F. nucleatum* subspecies *animalis*. Indeed, SimPlot analysis showed that the hypothetical new species/clade C1 displays 95% similarity to both *F. nucleatum animalis* and to *F. nucleatum vincentii* (Figure 5A). Most likely, the 95% identity between the new species/clade C1 and *F. nucleatum animalis* (clade C2) is higher than that calculated by Zepeda-Rivera and coworkers (92-93%) because we used only core genes. SimPlot analysis also confirmed the designation of taxonomic levels as species or subspecies to be problematic^11,12^. In fact, the core genomes of some species (e.g., *F. hwasookii* and *F. nucleatum polymorphum* or *F. nucleatum nucleatum* and *F. canifelinum*) were more closely related to each other than subspecies are among themselves.

### Population structure of the *F. nucleatum/F. periodonticum* lineage

To gain further insight into the population structure of fusobacteria, we used the program STRUCTURE, which relies on a Bayesian statistical model for clustering genotypes into populations, without prior information on their genetic relatedness^23–25^. The program can identify distinct subpopulations (or clusters, K) that compose the overall population. Subpopulations can then be related to specific features such as origin, classification, or phenotype. STRUCTURE is ideally suited to study highly recombining populations^23,26^.

Initially, we used the no admixture model, in which each individual is assumed to have derived from one of the modern populations. To estimate the optimal number of subpopulations in the *Fusobacterium* dataset, STRUCTURE was run for values of K from 1 to 12. The ΔK method yielded two peaks at K=2 and K=4 (Supplementary Figure 2). At K=2, STRUCTURE clearly separated the two main sub-lineages (*F. periodonticum/pseudoperiodonticum* and *F. nucleatum* plus related species) (Figure 6). At K=4, the four subpopulations paralleled the clusters identified in the PCA, and the new species/clade C1 was assigned to the population that includes *F. nucleatum vincentii* and *F. simiae,* not *F. nucleatum animalis* (Figure 6).

**Figure 6.**
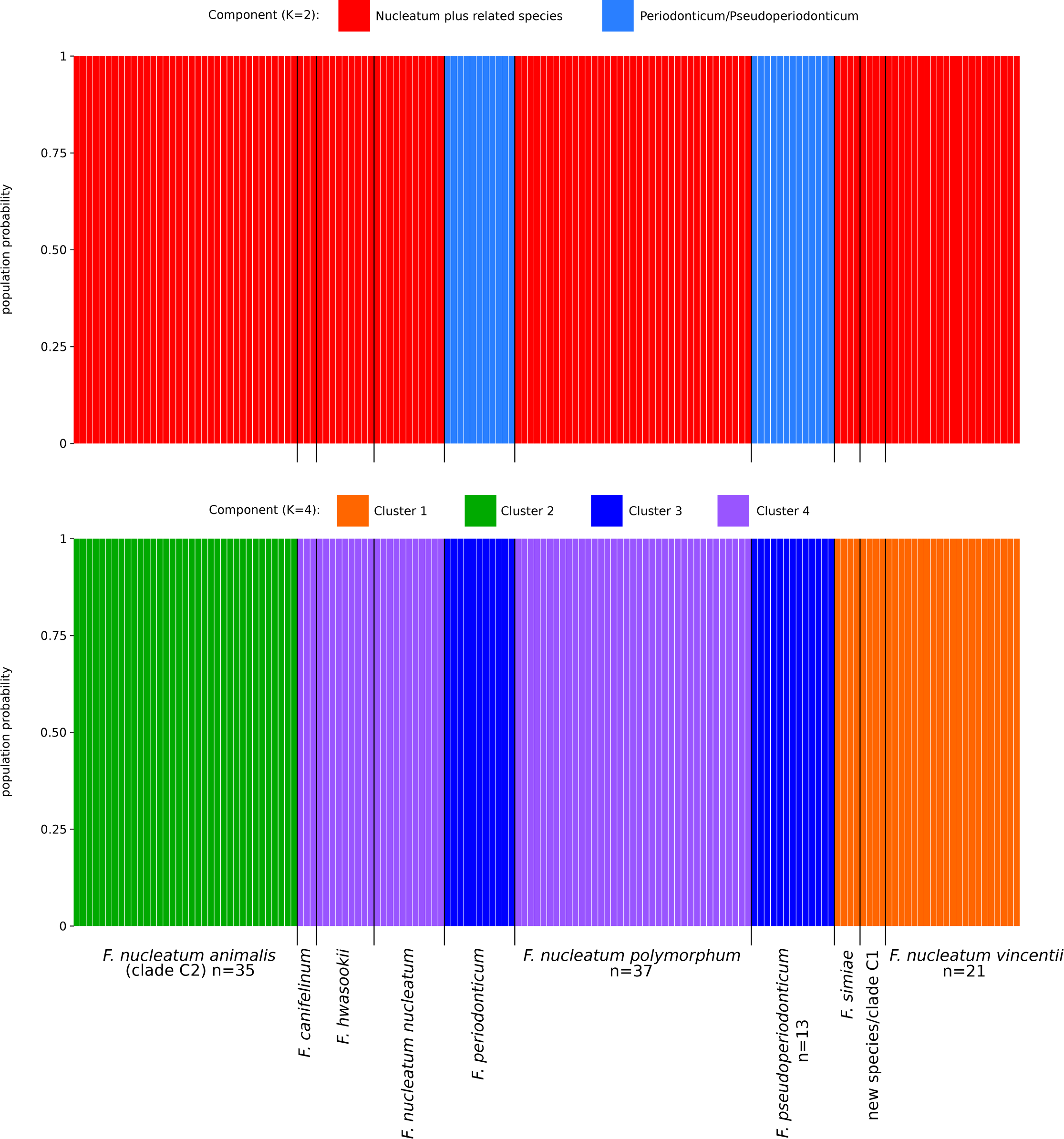
Population structure analysis: no admixture model. Bar plot representing the probability of population assignment from the STRUCTURE no admixture model. Each vertical line represents a *Fusobacterium* core genome. Results are shown for K= 2 and K=4. For the latter, populations are colored as the cluster in PCA analysis (Figure 5A).

In order to gain further insight into the evolutionary history of *Fusobacterium* core genomes, we repeated STRUCTURE analysis using the linkage model with correlated allele frequencies. This model assumes that discrete genome “chunks” were inherited from K ancestral populations^24^. The ΔK method identified two major peaks at K=3 and K=9 (Supplementary Figure 3). We thus analyzed in detail the results at K=9, which represents the finest level of structure for these genomes. Analysis of ancestry components showed that one of the ancestral populations (P_common) contributed variable proportions of ancestry to most genomes (Figure 7A). Other than this, individual ancestral components accounted most of the ancestry of distinct species or subspecies, with the only exception of *F. canifelinum,* which received ancestry components from 4 populations. Genomes of the new hypothetical species/clade C1 had most of their ancestry accounted by one of the nine ancestral populations, confirming they represent an entity distinct from *F. nucleatum animalis* (Figure 7A).

**Figure 7.**
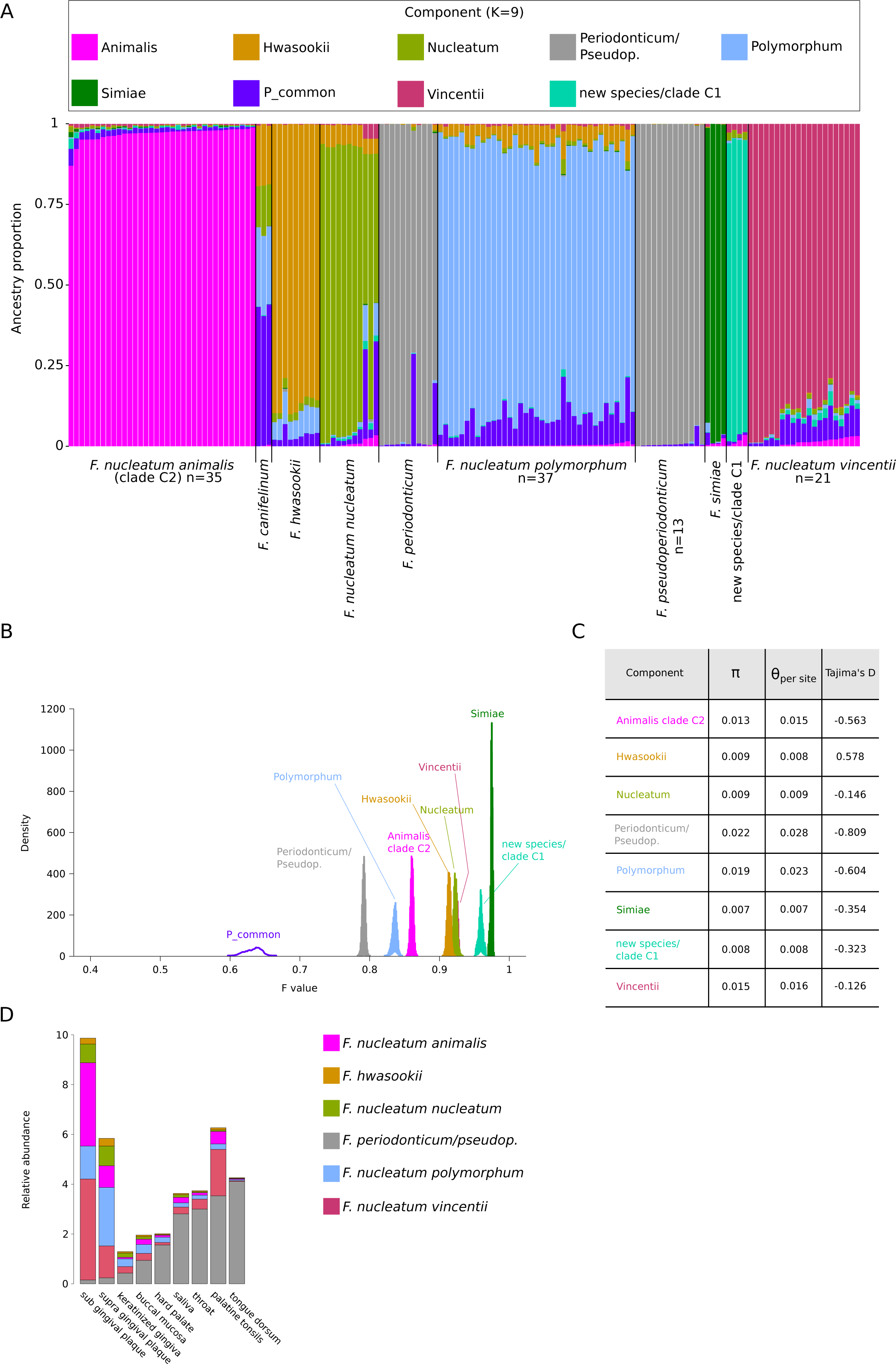
Population structure analysis: linkage model. **(A)** Bar plot representing the proportion of ancestral population components for K=9. Each vertical line represents a *Fusobacterium* core genome and it is colored by the proportion of sites that have been assigned to the nine populations by STRUCTURE. Ancestry components are named based on the genomes where they are more prevalent. **(B)** Distributions of F values for the nine populations. Colors are as in panel A. **(C)** Nucleotide diversity and Tajima’s for genomes that acquired a major part of their ancestry (>80%) from individual populations. **(D)** Relative abundance of different *Fusobacterium* species in different oral sites (data were derived from the expanded Human Oral Microbiome Database).

The linkage model also allows estimation of the F parameter, which represents a measure of genetic differentiation between populations based on allele frequencies. F can be interpreted as a measure of drift from a hypothetical common ancestral population. The lowest drift was detected for P_common, which is shared among most genomes (Figure 7B). However, the second population showing lowest drift was the one accounting for most ancestry of *F. periodonticum* and *F. pseudoperiodonticum.* The highest drift was instead obtained for the populations contributing ancestry to *F. simiae* and to the hypothetical new species/clade C1 (Figure 7B).

We next calculated nucleotide diversity and Tajima’s D for genomes that acquired a major part of their ancestry (>80%) from a single population. In line with the F results, the highest diversity was observed for the *F. periodonticum/F. pseudoperiodonticum* population, which also displayed the most negative value of Tajima’s D (Figure 7C). Overall, this is suggestive of a genetically diverse population that may have expanded in size. Conversely, low diversity was observed for *F. simiae* and the hypothetical new species/clade C1, which both showed moderately negative Tajima’s D, possibly suggesting that these populations have expanded after a bottleneck (Figure 7C).

Finally, we exploited the expanded Human Oral Microbiome Database (eHOMD) to compare the distributions of *Fusobacterium* species/subspecies among sites in the oral cavity and pharynx^27^. Data were available for six species/subspecies and indicated that *F. hwasookii* and *F. nucleatum nucleatum* are the least abundant across sites (Figure 7D). *F. nucleatum vincentii*, *F. nucleatum polymorphum* and *F. nucleatum animalis* display higher relative abundance but they are mostly specialized for the gingival plaque niche. Conversely, *F. periodonticum* seems to occupy different sites, including hard palate, tongue dorsum, palatine tonsils, throat, and saliva (Figure 7D). These results may be consistent with the idea that the *F. periodonticum/pseudoperiodonticum* population is more generalist in terms of distribution, whereas the other four species/subspecies might have drifted from the ancestral population as a consequence of niche adaptation.

## Discussion

Fusobacteria have attracted increasing interest in recent years due to their association with different human pathologies. Whereas *F. nucleatum* has been mainly investigated for its role in CRC, other *Fusobacterium* species were shown to contribute to different diseases. Thus, *F. necrophorum* causes tonsillitis and is commonly associated with Lemierre’s syndrome^28,29^, *F. periodonticum* was reported to contribute to oral squamous cell carcinoma^30^, *F. varium* was found in the guts of patients suffering from ulcerative colitis^31^, and *F. ulcerans* was associated with tropical ulcers^32^. Also, both *F. nucleatum* and *F. necrophorum* were isolated from liver abscesses and from cases of osteomyelitis or endocarditis^1,33–37^, and the same two *Fusobacterium* species are commonly found in the mucosal lesions of patients with appendicitis^38^. In addition to its role in CRC, *F. nucleatum* has been involved in other cancer types (bladder, oral, head and neck, cervical, and gastric) and in periodontitis^1,39^. Finally, *F. nucleatum* can colonize the placenta and cause preterm birth, intra-amniotic infection, stillbirth, neonatal sepsis, and hypertensive disorders of pregnancy^1,40,41^.

Because fusobacteria potentially contribute a huge health burden to human populations, molecular profiling approaches are essential to understand the epidemiology of fusobacteria-associated disease and to relate taxonomic groups to specific conditions. Even in the case of CRC, uncertainty still exists about which *Fusobacterium* species are pathogenic. A recent study showed that a lineage that includes *F. nucleatum* (all subspecies), *F. hwasookii*, *F. periodonticum,* and related fusobacteria is enriched in tumor samples and feces from CRC patients. A different lineage (represented by *F. varium* and *F. ulcerans)* was instead associated with lymphovascular invasion^9^. Conversely, another study found enrichment of *F. varium* in CRC samples^8^, whereas *F. nucleatum animalis* or even a specific clade within the diversity of this subspecies was found to be overabundant in the CRC niche in different studies^5,6,42^. Finally, *F. necrophorum* and *F. nucleatum* were found to colonize both primary CR tumors and the corresponding liver metastases^3^. Compared to CRC, other fusobacteria-related diseases have been investigated in shallower details and most commonly used approaches do not allow fine taxonomic definition. Moreover, as previously reported, we found that a number of *Fusobacterium* sequences in public repositories are miss-classified and several undescribed species exist, which complicates association analyses. We thus reasoned that a comprehensive investigation of the genetic diversity and relationships within the *Fusobacterium* genus might provide valuable information for future investigation to contextualize disease associations.

We first asked if and how intensely fusobacteria recombine, and whether recombination can affect phylogenetic relationships. Indeed, one of the effects of recombination is to unlink loci along the genome, so that they evolve independently and display diverse evolutionary histories. We report both ancestral (occurring during speciation) and recent (occurring after speciation) recombination between species in *rpoB*, which is commonly used as a marker gene and was proposed by Bi and co-workers for sample profiling^9^. As a consequence, different gene regions show distinct patterns of sequence similarity among species. In general, we found evidence of recent and ancestral recombination in all core genes we analyzed, indicating that fusobacteria have mosaic genomes and emphasizing that species/subspecies identification should not rely on single genes.

For a more detailed analysis, we focused on bacterial genomes in lineage 12 (from *rpoB* analysis), which show high sequence similarity in the analysis of core genes, as well. Although classified in different species and subspecies, these genomes are more closely related than most other fusobacteria and their relatively limited genetic diversity allows application of strategies to study population structure. Moreover, lineage 12 includes *Fusobacterium* species that have been intensively investigated for their role in CRC.

Both the PCA and STRUCTURE analyses were consistent in showing that modern populations are divided into two sub-lineages, which comprise *F. periodonticum/pseudoperiodonticum* and *F. nucleatum* plus related species. Further grouping was however observed, with four major populations defined as clusters in the PCA and as modern sub-populations in the non admixture STRUCTURE model. As is the case of genomes in the wider collection of the *Fusobacterium* genus, miss-classification or incomplete taxomomic definition was common within this lineage. However, using PCA or STRUCTURE analysis most genomes could be assigned to known species or subspecies.

Nonetheless, all analyses confirmed that four genomes that include strains FNU and 13-08-02 (in clade C1 in Zepeda-Rivera et al.) do not belong to the *F. nucleatum animalis* subspecies or any other known species/subspecies. Thus, our data do not support the previously suggested division of *F. nucleatum animalis* diversity in two clades^6^. In their recent study, Zepeda-Rivera and coworkers showed that clade C2 was associated to the CRC niche, whereas C1 was not. The two clades were suggested to differ in terms of accessory genome size, number of extrachromosomal plasmids and immune defences, as well as methylation patterns, representation of adhesins, and metabolic potential. The cells of bacteria in clade C2 were also found to be longer and thinner than those in clade C1, and to have a higher level of cancer cell invasion. All these differences would be noteworthy for bacteria in the same species, but we show here that this is not the case. Indeed, all these features were compared between two distinct species, although closely related. Whereas this is unlikely to change the conclusion that *F. nucleatum animalis* is associated with CRC, our data call for a re-assessment of the characteristics of clade C2, which were established in comparison to a different species^6^. This is particularly true in light of the PCA and STRUCTURE analyses, which indicated that the new specie/clade C1 and *F. nucleatum animalis* belong to different clusters or modern populations. Notably, the linkage model STRUCTURE analysis showed that the new species/clade C1 inherited ancestry from a distinct ancestral population that experienced substantial drift in comparison to most other populations, including the one that contributed to the ancestry of *F. nucleatum animalis*. Genetic drift was shown to promote genome reduction and decreased coding density in bacteria^43^, in line with the small genomes of clade C1 bacteria^6^. The new species/clade C1 also shows very limited genetic diversity, although analysis is not particularly robust, as it was necessarily limited to four genomes. Overall, these results indicate that more extensive and lineage-wise comparisons are necessary to establish which *F. nucleatum animalis* characteristics contribute to CRC association.

Interestingly, STRUCTURE results showed that the ancestral population from which *F. periodonticum/pseudoperiodonticum* emerged experienced the lowest drift and these species now display the highest genetic diversity. Together with the negative Tajima’s D, this is suggestive of a size expansion in the population. Compared to other oral *Fusobacterium* species, whose populations experienced stronger drift, *F. periodonticum* was found to commonly occur in different oral sites, with lower representation in the gingival plaque. Conversely, *F. nucleatum* subspecies were mostly specialized for plaque. These results are consistent with the view that, during their evolution *F. nucleatum* subspecies drifted away from a common ancestral population to colonize a new niche. Our work has limitations. One of the most serious concerns the scant meta-data available for the *Fusobacterium* genomes we analyzed. For most of them, we had no information about the origin in terms of geographic location, body site, or host. In most cases, the health status of the host was unknown, as well as the isolation source. Whereas this limits the possibility to perform more detailed analysis of genetic diversity in fusobacteria, the inability to control for external variables might also introduce unrecognized biases. For instance, individual bacterial species were shown to be more genetically diverse among African than non-African human hosts^26,44–47^. The unequal representation of genomes from different geographic areas in different species might thus affect our measures of nucleotide diversity. Another limitation concerns our focus on core genomes, which was motivated by the need to obtain reliable alignments and PI sites, as well to maintain data tractable for STRUCTURE analysis. Although, bacterial pangenomes are known to be highly diverse and virulence factors are often encoded by accessory genes, the purpose of our work was to describe the genetic relationships in the *Fusobacterium* genus. We consider that these data might be valuable to develop a much needed molecular profiling approach that can shed light into the epidemiology of fusobacteria-associated diseases.

## Methods

### Bacterial and core genes sequences

The list of *Fusobacterium* genomes was derived from the BV-BRC site (https://www.bv-brc.org/, as of July 2023) by selecting entries with "good" genome quality. Complete and draft genome sequences were obtained from the NCBI database by using the getGenome function from the R package biomaRt^48^; the set consisted of 361 bacterial samples (Supplementary Table 1). Complete and metagenome-assembled genomes were used as input data for the Genome Taxonomy Database Toolkit (GTDB-Tk)^49^. This tool provides an automated taxonomic classification of bacterial sequences based on a set of 120 single copy marker proteins; GTDB-Tk also identifies and extracts from input genomes both nucleotide and protein sequences of each marker. The nucleotide sequences were then used for the subsequent analyses. Since not all samples have the whole genome covered, we were unable to retrieve all markers for all samples.

### *rpoB* gene alignment and network

An alignment based on 345 *rpoB* gene sequences was constructed using MAFFT with default parameters^50^. A neighbor-net split network was generated throughout SplitsTree4^51^: a data matrix was generated from the aligned sequences, estimating distances with the HKY85 model and removing parsimony-uninformative and constant sites.

### Recombination and sequence similarity analyses

The same *rpoB* alignment described above was used to run fastGEAR, an algorithm that detects recombination events between inferred lineages, as well as from external origins. In particular, this method first clusters sequences into lineages, then it identifies both recent (i.e. affecting a subset of strains in a lineage), and ancestral (i.e. affecting all strains in a lineage) recombination events^52^. The same approach was used to identify ancestral and recent recombination events for a list of 45 genes from the 120 marker genes. These 45 genes were selected because they were longer than 1000 nucleotides and they were present in at least 300 genomes (Supplementary Table 1). The FastGEAR output was then used to generate a plot of recent recombination events versus ancestral ones, normalized by gene alignment length.

A concatenated alignment, based on 120 core genes, was generated with the same genomes used in the *rpoB* alignment. The alignment was generated using the GUIDANCE2 suite^53^, setting sequence type as nucleotides and using MAFFT as an aligner. GUIDANCE2 also allows to filter unreliably aligned positions. We thus removed positions with a score lower than 0.90^54^.

Sequence similarity analyses were performed using SimPlot++^55^. This tool generates a similarity network plot based on a multisequence alignment. Each node of the network represents a sequence or group of sequences and edges indicate the global (over the whole sequence) or local (over one or more sub-regions) similarity among nodes.

### PCA, population STRUCTURE, and nucleotide diversity

Strains belonging to lineage 12 were selected to build a new concatenated gene alignment. Concatenated gene sequences that were shorter than 80% of the longest sequence were discarded from the analyses: this filtering allowed us to limit the number of gaps in the alignment but also to take into account differences in gene lengths; this generated a set of 148 strains. We then generated an alignment by applying GUIDANCE2 as described above. From this new alignment, biallelic parsimony-informative (PI) sites were extracted; in particular, we selected biallelic sites, each with a minimum frequency of two, for those genomic positions where at least 50% of sequences had non-missing information. Gaps and all nonstandard nucleotide bases were considered as missing values. This generated a list of 26430 variable positions. Principal component analysis (PCA) was performed with the mixOmics R package^56^, using the PI matrix as input and two principal components. The same PI data was also used to run STRUCTURE. First, the software was run with K=1 to estimate the frequency spectrum parameter (λ), as suggested^24^. The λ parameter was estimated to be equal to 0.5878. Using this value, both the no admixture model with independent allele frequencies and the linkage model with correlated allele frequencies were run^23,24^. Both models were run with different values of K populations, from 1 to 12. In particular, for each K, ten runs with a MCMC total chain length of 500,000 iterations and 50,000 iterations as burn-in were run. The optimal K was evaluated with Evanno’s method^57^ using the HARVESTER tool^58^. The CLUMPAK^59^ software was used to combine replicate runs from the same K and to generate the Q value matrix. For the linkage model analysis, the amount of drift that each subpopulation experienced from a hypothetical ancestral population was quantified by the F parameter calculated for the optimal k value^24^.

Finally, results obtained from the linkage model were used to group strains to estimate population genetic parameters. Specifically, each strain was assigned to one of the defined K populations if it had an ancestry component higher than 80% for that specific population (i.e. admixed individuals were excluded); then nucleotide diversity and Tajima’s D were calculated for each populations using the DnaSP software^60^.

### Relative abundance data

Bacterial abundance in the human mouth and aerodigestive tract was retrieved from the Human Oral Microbiome Database v3.1 (https://www.homd.org/)^61^. We retrieved the relative abundance of *Fusobacterium* species/subspecies from three different experiments and we calculated mean values for 9 different oral districts: buccal mucosa, keratinized gingiva, hard palate, tongue dorsum, palatine tonsils, throat, saliva, supra-gingival plaque, and sub-gingival plaque.

## Acknowledgments

This work was supported by the Italian Ministry of Health (“Ricerca Corrente” to MS).

## Author Contributions

Conceptualization, M.S and D.F.; Methodology, M.S., C.M., and D.F.; Investigation, C.M., D.F., R.C.; Writing Original Draft, M.S. C.M.; Writing Review & Editing, M.S., D.F., and R.C.; Funding Acquisition, M.S.

