## Supplemental figures 1-3, Supplemental Table 2 for "Tangled genetic relationships within the *Fusobacterium* genus"

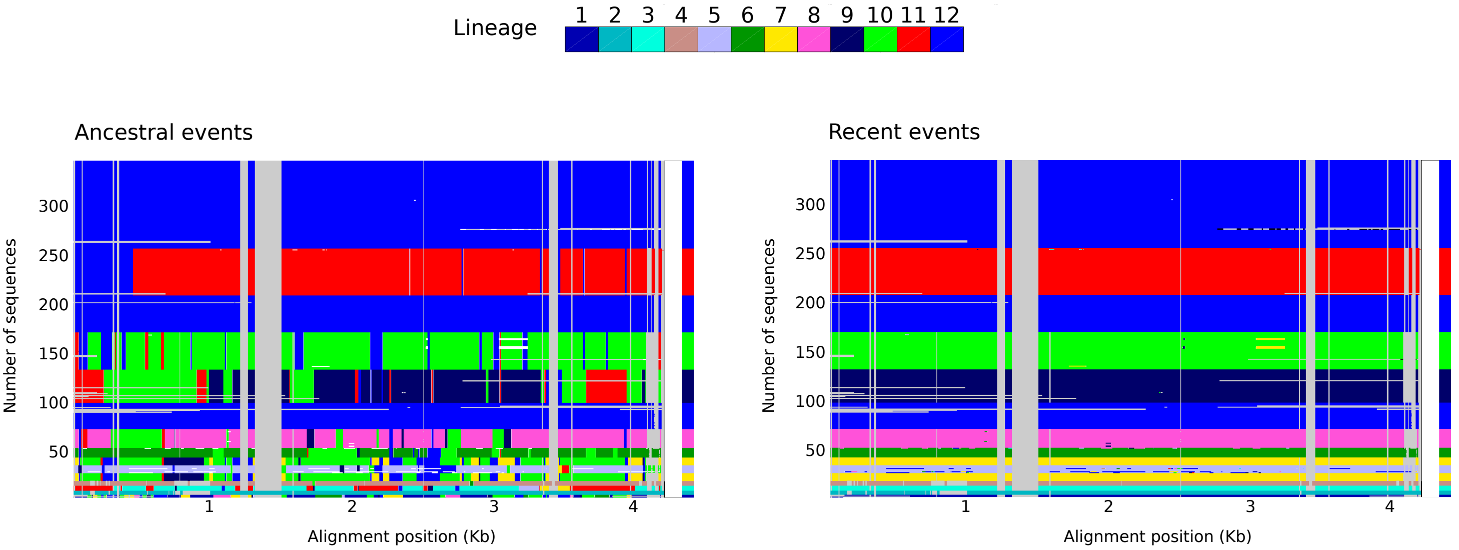


**Supplementary Figure 1. Plot of fastGEAR results on the rpoB alignment.** Ancestral recombination events are plotted on the left, recent ones on the right. Recombination between lineages is depicted with different colors. Gray columns represents gaps in the alignment. For the ancestral recombination plot, recent recombination events are shown as white gaps.

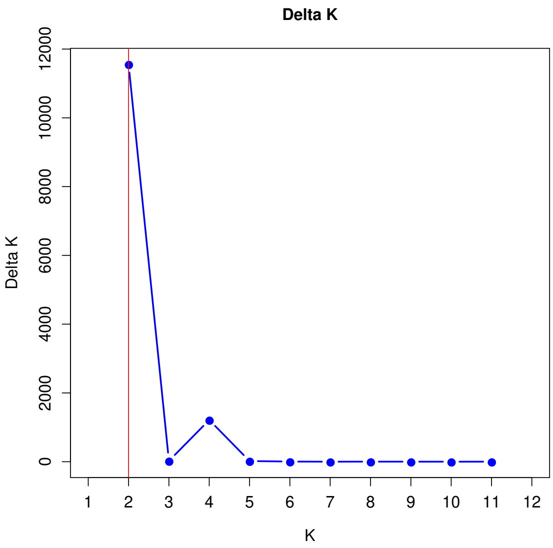


**Supplementary Figure 2. Analysis of optimal K for the STRUCTURE no admixture model.** ΔK is calculated as ΔK = mean(|L''(K)|) / sd(L(K)). The peaks of this distribution are the optimal K used in STRUCTURE analysis.

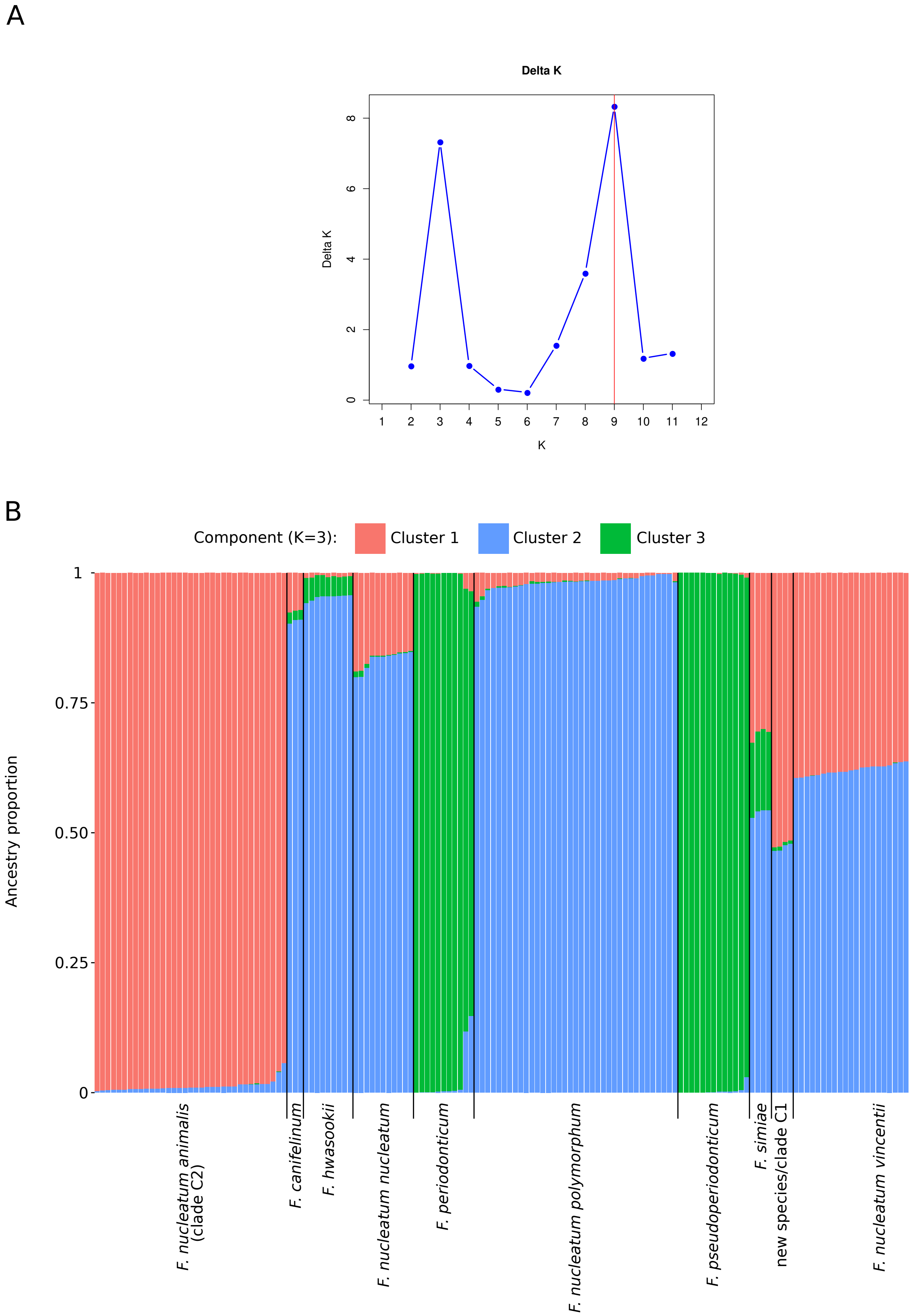


**Supplementary Figure 3. Analysis of STRUCTURE linkage model.** (A) Evanno's method plot for optimal K. ΔK is calculated as ΔK = mean(|L''(K)|) / sd(L(K)). The peaks of this distribution are the optimal K used in STRUCTURE analysis. (B) Bar plot representing the proportion of ancestral population components for K=3. Each vertical line represents a Fusobacterium core genome and it is colored by the proportion of sites that have been assigned to one of the populations by STRUCTURE.

**Supplementary table 2. List of recombination events identified by fastGEAR.**

| Gene | Number of Strains | Number of Recent Events | Number of Ancestral Events |
|---|---|---|---|
| *prfA* | 348 | 14 | 37 |
| *prfB* | 345 | 50 | 93 |
| *rseP* | 352 | 70 | 59 |
| *ftsZ* | 346 | 53 | 49 |
| *ychF* | 358 | 36 | 73 |
| *tig* | 350 | 99 | 91 |
| *uvrC* | 342 | 88 | 231 |
| *alaS* | 355 | 67 | 143 |
| *ileS* | 312 | 70 | 135 |
| *leuS* | 353 | 192 | 153 |
| *metG* | 348 | 49 | 192 |
| *serS* | 355 | 33 | 89 |
| *radA* | 355 | 135 | 109 |
| *cysS* | 352 | 28 | 152 |
| *hisS* | 351 | 38 | 92 |
| *argS* | 346 | 65 | 107 |
| *aspS* | 354 | 69 | 118 |
| *pheT* | 350 | 113 | 215 |
| *infB* | 355 | 41 | 246 |
| *hemN* | 338 | 69 | 148 |
| *mfd* | 355 | 400 | 446 |
| *polA* | 338 | 374 | 405 |
| *uvrB* | 353 | 164 | 259 |
| *recN* | 347 | 159 | 114 |
| *recG* | 354 | 190 | 286 |
| *rpsA* | 345 | 103 | 62 |
| *purB* | 347 | 59 | 42 |
| *ffh* | 348 | 53 | 80 |
| *secA* | 348 | 122 | 245 |
| *secY* | 342 | 12 | 40 |
| *atpD* | 348 | 31 | 66 |
| *gyrB* | 341 | 79 | 37 |
| *gyrA* | 346 | 92 | 64 |
| *murC* | 352 | 71 | 88 |
| *murD* | 352 | 165 | 174 |

| | | | |
|---|---|---|---|
| *guaB* | 349 | 21 | 51 |
| *dnaG* | 343 | 183 | 195 |
| *lepA* | 343 | 28 | 67 |
| *typA* | 349 | 40 | 52 |
| *nusA* | 352 | 52 | 76 |
| *rpoB* | 345 | 98 | 198 |
| *dnaK* | 350 | 26 | 60 |
| *rpoC* | 348 | 94 | 235 |
| *dnaX* | 346 | 298 | 187 |
| *der* | 346 | 32 | 71 |
